# Modelling complex growth profiles of *Bacteroides fragilis* and *Escherichia coli* on various carbohydrates in an anaerobic environment

**DOI:** 10.1101/2023.05.01.538938

**Authors:** Zachary McGuire, Sainitya Revuru, Sheng Zhang, Amanda Blankenberger, Moiz Rasheed, Jacob D. Hosen, Guang Lin, Mohit S. Verma

**Author notes:** Equal contribution (Zachary McGuire, Sainitya Revuru, Sheng Zhang, Amanda Blankenberger).

## Abstract

Previously published models for microbial growth focus only on either death or growth and are unable to account for differently shaped growth curves. Currently, there is no model capable of incorporating combinations of microbial growth trends. This study creates a bacterial growth model that incorporates growth, death, lag, and tail phases as well as applies this model to the growth trends of *Bacteroides fragilis* and *Escherichia coli* on 13 different carbohydrate substances. Growth trends were collected by measuring the optical densities over 72 hours for either *B. fragilis* or *E. coli* in a chemically defined media supplemented by a mono- or disaccharide. The Digital Environment to Enable Data-driven Science (DEEDS) platform was utilized to parse data and apply the developed model to obtain parameter values. *E. coli* was found to grow on the chemically defined media alone while *B. fragilis* was unable to grow on it alone. *E. coli* growth was led by 10 mM concentration of substrates while *B. fragilis growth* was substrate dependent. Bacterial death only occurred for *B. fragilis* but was found to be dependent on concentration for the two most significant substrates. A singular model was developed that does not require prior knowledge of metabolomics and is capable of incorporating a combination of growth and death trends.

## 1. Introduction

The most widely used models for bacterial growth predict sigmoidal-shaped curves, which do not represent the entirety of the growth data observed in a closed system (since these include a death phase that is often ignored in growth models). In order to quantify a growth curve and perform statistical analysis on the influence of environmental conditions (e.g., different carbon sources), we need a model that can fit both growth and death phases without segmenting the data. Here, we present such a model by introducing a death rate term to the logistic growth model and use this model to quantify growth curves with and without death phases. We use this model to obtain and evaluate growth and death parameters in the presence of 13 different carbon sources at three concentrations each for two different bacteria: *Bacteroides fragilis* and *Escherichia coli* as seen in the workflow demonstrated in Fig. 1.

**Figure 1.**
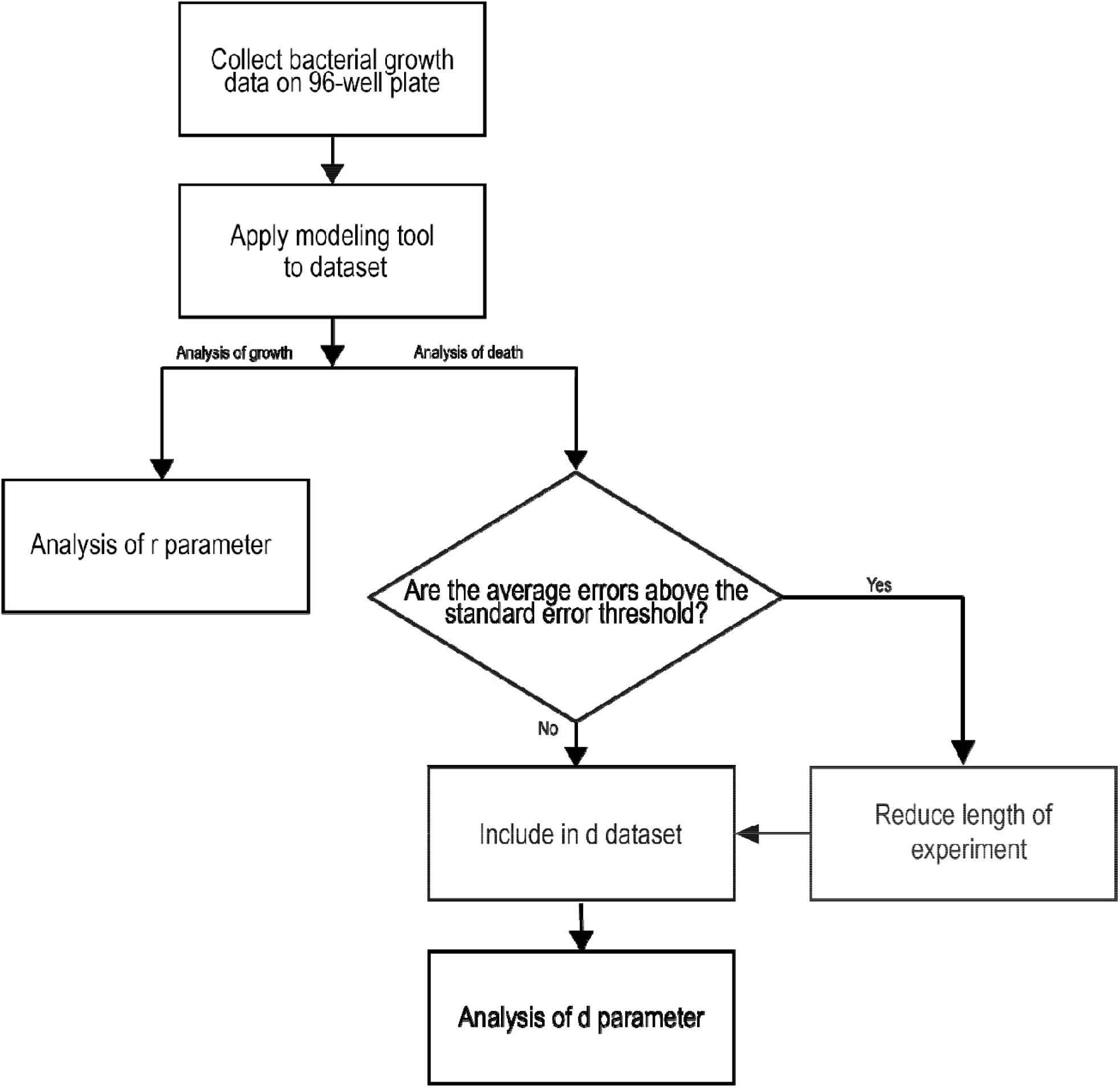
Schematic of experimental setup including the logic for including values in death parameter analysis. r is the instantaneous growth rate. d is the linear rate of decline of carrying capacity.

Published models in predictive microbiology tend to focus on either growth or death depending on the intended use in food safety and quality. Growth models focus on studies in food hazards and spoilage^1^, while death models focus on attempts to kill bacteria in efforts to improve shelf life and food safety^2^. The death phase of the bacterial growth curve that occurs after bacteria reach their carrying capacity has largely been ignored. Since most of the work in this field has applications in food microbiology, the death phase may have been ignored in modelling as food cannot be unspoiled. Furthermore, not all experimental bacterial growth data will show a death phase, depending on the length of the experiment and the environmental conditions. The stationary phase of bacterial growth can be maintained for extended periods of time. The amount of time spent in each phase of the bacterial growth curve is dependent on the species of the bacteria as well as the environment, so the length of the stationary phase or the time until rapid death will occur is difficult to predict *a priori*^3^.

Many models for non-sigmoidal growth require additional inputs which may not be available to the researcher or possible to obtain given the bacterium of interest. Published literature has the following three limitations: i) cybernetic models require growth studies of individual substrates in minimal media^4^ (which prohibits growth of certain bacteria because they may not have the necessary micronutrients in minimal media), ii) diauxic models for biphasic growth require knowledge of enzyme kinetics^5^, iii) many models are designed for specific trends observed in growth, e.g., sigmoidal^6,7^, biphasic^4,5^, death^2,7^, death then recovery growth^7^. A comparison of the published models with the model in this study is shown in Table 1. ^8 9 10 11^

**Table 1.**
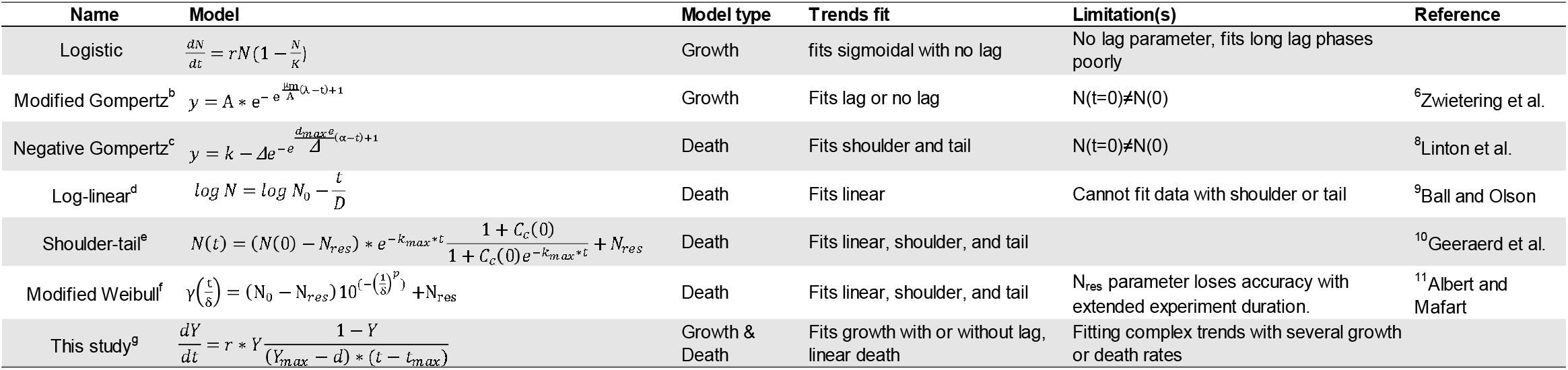
Comparison of growth and death models with the model of this study. ^a^ N = instantaneous population size, r = instantaneous growth rate, and K = carrying capacity. ^b^ λ = lag time, µ_m_ = maximum specific growth rate, A = carrying capacity. ^c^ k = initial cell count, Δ = decrease of cell count over time, d_max_ = maximum death rate, α = shoulder. ^d^ D = decimal reduction value, N and N_0_ = population at time t. ^e^ K_max_ = maximum death rate, N_res_ = residual population, C_c_ = concentration of the critical component. ^f^ γ(t) = bacterial concentration at time t. p = curve shape (concavity, convexity, and shoulder). t = time, δ = first decimal reduction concentration for the population not belonging to N_res_, N(0) = initial cell concentration, N_res_ = residual bacterial concentration. ^g^ r = instantaneous growth rate, Y = instantaneous population size, Y_max_ = max population size, d = linear rate of decline of carrying capacity, t_max_ = time at which Y_max_ is reached. In all cases, t = time.

To our knowledge there are limited models that can be used to model any combination of the trends listed above. A model is necessary that is versatile and can fit many different trends in bacterial growth while providing meaningful parameters that can be statistically analyzed. Here, we present a growth model that can fit variable growth profiles, both with and without a death phase. We apply this model to several different trends observed in the growth of *Bacteroides fragilis* and *Escherichia coli* on 13 different substrates to demonstrate the versatility of the model in fitting many different plot shapes.

## 2. Materials and Methods

### 2.1 Bacterial strains and culture

The bacterial strains used in this study include *Bacteroides fragilis* (ATCC 43858) and *Escherichia coli* (LF82 strain obtained from Professor Ramnik Xavier’s Lab). Frozen secondary stocks of the bacteria were prepared (1 mL of culture and 1 mL of 50% glycerol in a 2 mL cryovial) and stored in -80 °C. *B. fragilis* is cultured in commercially available chopped meat broth (Pre-Reduced II Hungate Cap, BD BBL™) and *E. coli* LF82 is cultured in Tryptic Soy Broth (BD Biosciences). Both the bacteria are incubated at 37 °C and allowed to grow under anaerobic conditions for two days with a gas composition of 4% H_2_, 5% CO_2_, and 91% N_2_. The cells are incubated in a Heated Vinyl Anaerobic Customized Type B Chamber with a hydrogen sulfide removal column (Coy Laboratory Products Inc., Grass Lake, MI).

After incubating for two days, the cultures are tightly sealed and centrifuged at 10,000 rpm for 5 minutes. The supernatant is discarded and the pellets are suspended in required volumes of phosphate-buffered saline (PBS, 137mM NaCl, 2.7mM KCl, 10 mM Na_2_HPO_4_, 1.8mM KH_2_PO_4_, pH 7.4) in anaerobic conditions to yield OD_600_ of ∼0.5 as measured in a Corning® 96-well clear flat bottom polystyrene microplate (Catalog #3370, Corning®, Corning, NY) by pipetting 250 µL and using a Biotek® Epoch™ 2 multi-well plate reader (Biotek Instruments, Winooski, VT).

### 2.2 Defined media preparation

In this study, the bacteria are allowed to grow in a modified chemically defined base medium ZMB1^12^ (Table S1) supplemented with different carbohydrates (Fig. 2). ZMB1 is chosen as the base medium because it supports high cell density growth of the selected bacteria^12^. Detailed modified ZMB1 preparation information can be found in Table S1.

**Figure 2.**
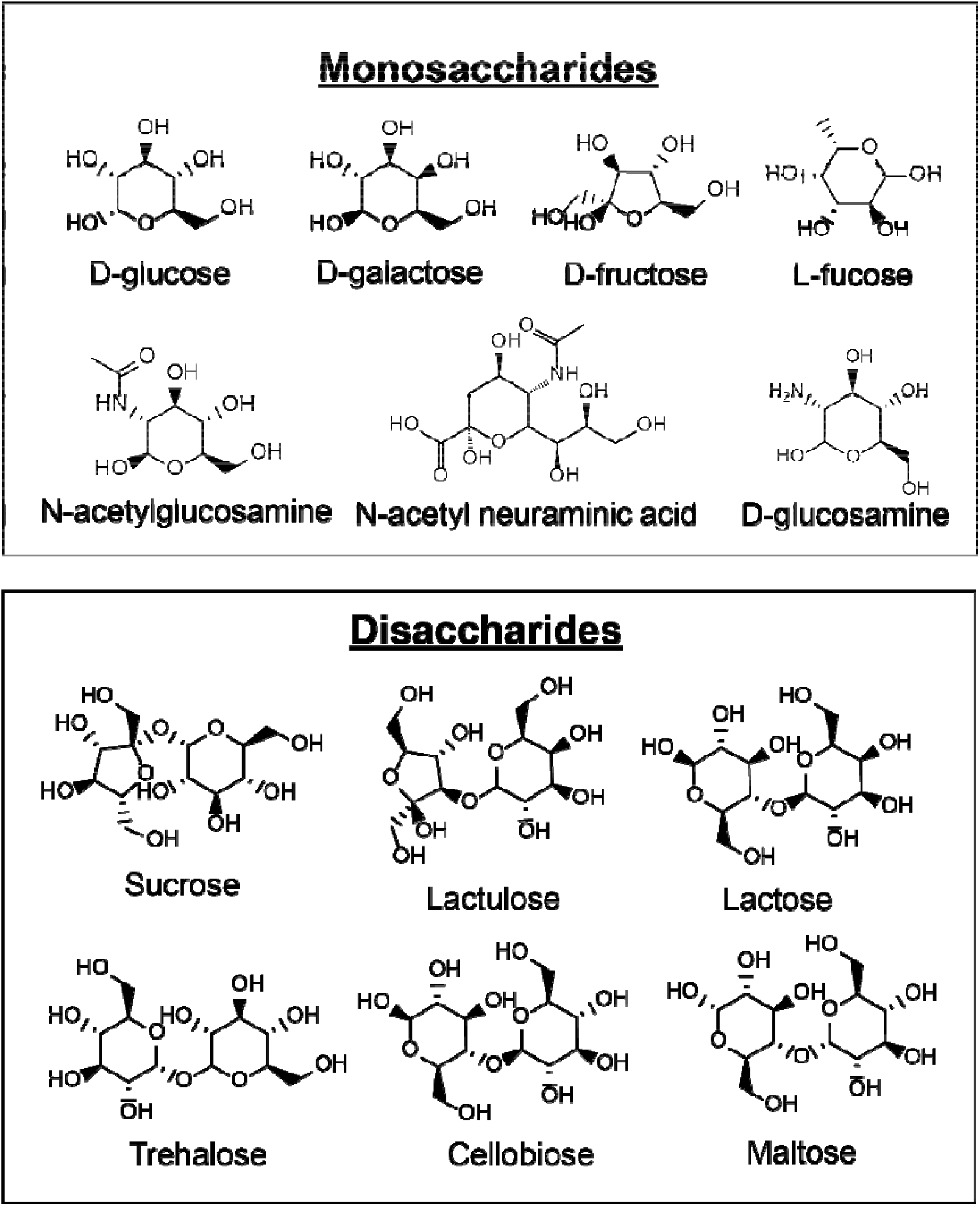
Chemical structures of the monosaccharides and disaccharides used in this work.

Carbohydrate (Fig. S1) stocks are prepared in concentrations of 111 mM as exemplified in Table S2. The solution is then filtered using a Steriflip® Vacuum-driven Filtration System (Catalog # SCGP00525, EMD Millipore Corporation, Burlington, MA). 111 mM carbohydrate stocks were then serially diluted using two 10x dilutions into sterile modified ZMB1 to create solutions at concentrations of 11.1 mM and 1.11 mM. If necessary, the pH of the solution is adjusted to a value of 7.0 using 10N NaOH.

### 2.3 Experimental design and analysis

The experiments are carried out in a Corning® 96-well clear flat bottom polystyrene microplate (Catalog #3370, Corning®, Corning, NY) to observe the growth of *B. fragilis* and *E. coli* LF82 in all 13 carbohydrates over time (72 hrs) separately. For every set of growth condition, eight technical replicates are used. 225 μL of prepared media (modified ZMB1 and carbohydrates) are added into the microtiter plates. 25 μL of the inoculum (bacteria in PBS) is then transferred into the wells containing the prepared media and covered with 50 μL sterile mineral oil (Catalog #S25439, Fisher Science Education, Nazareth, PA) to prevent evaporation. This dilution yields final experimental concentrations of 100 mM, 10 mM, and 1 mM for each condition. Two blanks (modified ZMB1 and PBS sans bacteria) and negative controls (bacteria in PBS and ZMB1 sans carbohydrates) of the same quantities are used for comparison (Fig. S2).

Optical density of the microplates is measured at 600 nm (OD_600_) over 72 hours at an interval of 30 mins using Epoch 2 Microplate Spectrophotometer (BioTek Instruments, Inc., Winooski, VT) with the Corning® 96-well clear flat bottom polystyrene microplate #3370. Bacterial growth is estimated via accumulation of biomass by subtracting the average of the blanks from each experimental condition from the average of the technical replicates for that same condition. The reading process for multiple plates is automated using a BioStack™ 4 microplate stacker (BioTek Instruments, Inc., Winooski, VT), which cycles the experimental 96-well microplates and removes the lids at the specified interval as to not interfere with spectrophotometric values. After the Epoch 2 Microplate Spectrophotometer is finished reading each plate, the BioStack™ 4 microplate stacker returns the lid to the microplate and places the microplate back into the stacker to wait for the next reading interval to begin.

### 2.4 Computational Model

According to the logistic model of population growth, we have

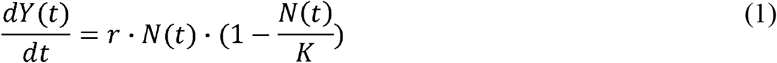

where Y is the population size, K is the carrying capacity, and r is the per capita growth rate. In our setting, the carrying capacity decreases over time as the resource is consumed. We use time-dependent carrying capacity as follows:

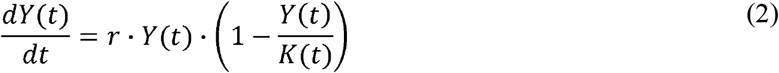

Denote t’ = argmax_t Y(t). Since 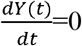 when t=t’, we have

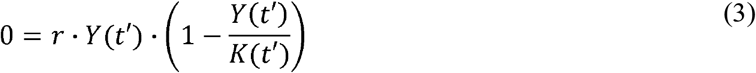

or

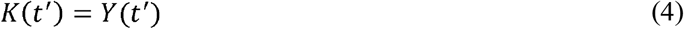

Suppose the carrying capacity decreases over time in the rate of d. In other words,

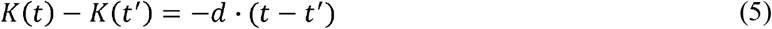

We have

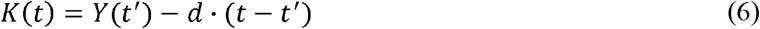

Substitute Eq. (5) into Eq. (2), and we have

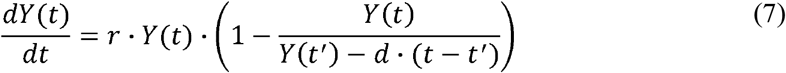

Given each dataset, we approximate dY(t)/dt numerically, and then obtain the growth rate and the carrying capacity decrease rate for the dataset by solving the following optimization problem:

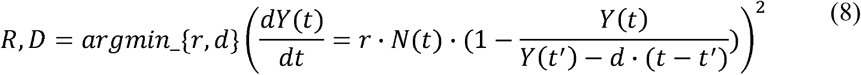

### 2.5 DEEDS Workflow

The platform Digital Environment for Enabling Data-Driven Science (DEEDS) was used to analyze the data presented in this paper. The raw data from the Gen5™ software (BioTek Instruments, Inc.) was uploaded here. Two codes were ran in series to generate the parameter data presented in this paper.

The first Python code (parsing.py) uses two files for input: the Gen5™ software’s Microsoft Excel document and a separate Microsoft Excel document detailing the experimental conditions of each well of each plate in an experiment. The code then parses, labels, and subtracts the carbohydrate blanks for all experimental conditions. The code creates a separate .csv file for each experimental condition on an experiment. See Supporting Information for the code.

The second Python code (modelFit.py) uses the parsed .csv files from the previous code to analyze the experimental data used in the model presented above. See Supporting Information for the code.

Since this work was initially completed, the DEEDS platform is no longer available or managed. Thus, we are providing the python code in supporting information to help replicate the results reported here.

## 3. Results and Discussion

### 3.1 Selection of carbohydrates for growth curves

A total of 13 carbohydrates are chosen to observe the growth of *B. fragilis* and *E. coli* LF82 in this study (Fig. 2A and B). D-glucose, a simple sugar, is a common carbon source for most bacteria. We wanted to analyze the growth of bacteria in both mono- and di-saccharides - 1) by including isomers of D-glucose (D-fructose and D-galactose), 2) by removing and adding functional groups (L-fucose, D-glucosamine, N-acetyl-D-glucosamine), 3) by including sugars that are a resultant of different glycosidic linkages between two glucose molecules and finally, 4) by including di-saccharides that are a resultant of linkages between two different monosaccharides (Fig. S1).

### 3.2 Optical density growth curves for *B. fragilis*

The growth of *B. fragilis* in ZMB1 media with individual carbohydrate sources varies per source and concentration of sugar. *B. fragilis* did not grow in ZMB1 without a sugar present. Like *E. coli, B. fragilis* has been reported to utilize amino acids for growth but in studies with media consisting of only salts, minerals, and amino acids there was only a small amount of initial growth^13^. Most notably, *B. fragilis* demonstrated a lag phase on most growth curves. It has been hypothesized previously that at CO_2_ levels like the 5% used in these experiments is not high enough for the bacteria to be able to grow^14^. So in the lag phase, the bacteria are creating more CO_2_ until a level is reached in which the bacteria are able to replicate^14^.

*B. fragilis* had significant growth rates on all sugars except trehalose and cellobiose. The bacteria has been previously identified by its inability to ferment trehalose^15^. Similarly, one of the defining differences among other *Bacteroides* species is *B. fragilis’* inability to ferment cellobiose^16^. Unlike *E. coli*, all *Bacteroides* species are capable of fermenting maltose^16^. Maltose fermentation utilizes the RokA kinase which is also used to phosphorylate hexose and amino sugars^17^.

In *B. fragilis*, N-acetyl glucosamine is used more efficiently than glucose and previously published results suggest similar findings^18^. We also found that neuraminic acid was utilized by *B. fragilis*. The kinase responsible for the catabolism of both neuraminic acid and n-acetyl glucosamine is RokA^17^.This kinase along with the hexA kinase are the only glucose kinases in the cell^17^.

### 3.3 Optical density growth curves for *E. coli* LF82

The growth of *E*.*coli* in ZMB1 also varied with the different carbohydrate sources. It should be noted that *E. coli* was able to grow in the modified ZMB1 without an added carbohydrate source. *E. coli* is typically grown on media containing Bacto™ Tryptone which is a supplier of amino acids for the bacteria^19,20^. *E. coli* has been known to grow on L-alanine as its sole carbon source^21^. It is probable the bacteria were able to utilize the amino acids included in the modified ZMB1.

*E. coli* demonstrated significant growth on all monosaccharides, but only showed limited growth on some disaccharides. Cellobiose and maltose both contain two glucose monomers but the bacteria had little to no growth. Other strains of *E. coli* have been able to utilize cellobiose through mutations likely not found in the LF82 strain used^22^. *E. coli*’s ability to utilize maltose such as in the K12 and O157:H7 strains rose from a mutant of the original strain^23,24^. When two glucose monomers are connected in an α-glycosidic linkage as in trehalose, there is significant *E. coli* growth. The K12 and LF82 strains are capable of fermenting trehalose^25^. Additionally, the growth curves of the 10 mM and 100 mM differ likely due to trehalose being degraded to glucose by different methods dependent on the osmolarity of the growth conditions^26^.

In previously published studies, *E. coli* is able to utilize N-acetyl-D-glucosamine more efficiently than glucosamine^27^. Glucosamine reports a higher growth rate in this study likely due to a mutation that limits *E. coli* growth on N-acetyl-D-glucosamine^28^. This mutation is likely on the glucosamine-6-phospahte deaminase (NagB) which when affected allosterically causes differences in growth rates^27^. *E. coli* is also capable of utilizing neuraminic acid well due to the presence of an extracellular mutarotase (NanM) in *E. coli* allowing neuraminic acid to be utilized as a carbohydrate source^29^.

### 3.4 Analysis of model parameters

The r and d model parameters show growth or death contribution, respectively, to the growth curve by an undetermined mechanism (Fig. 4 and 5). In this model, the r parameter is defined the same way as the logistic equation. The d parameter describes the speed of decline of the carrying capacity of the population over time. When d = 0, the model becomes the logistic model.

**Figure 3.**
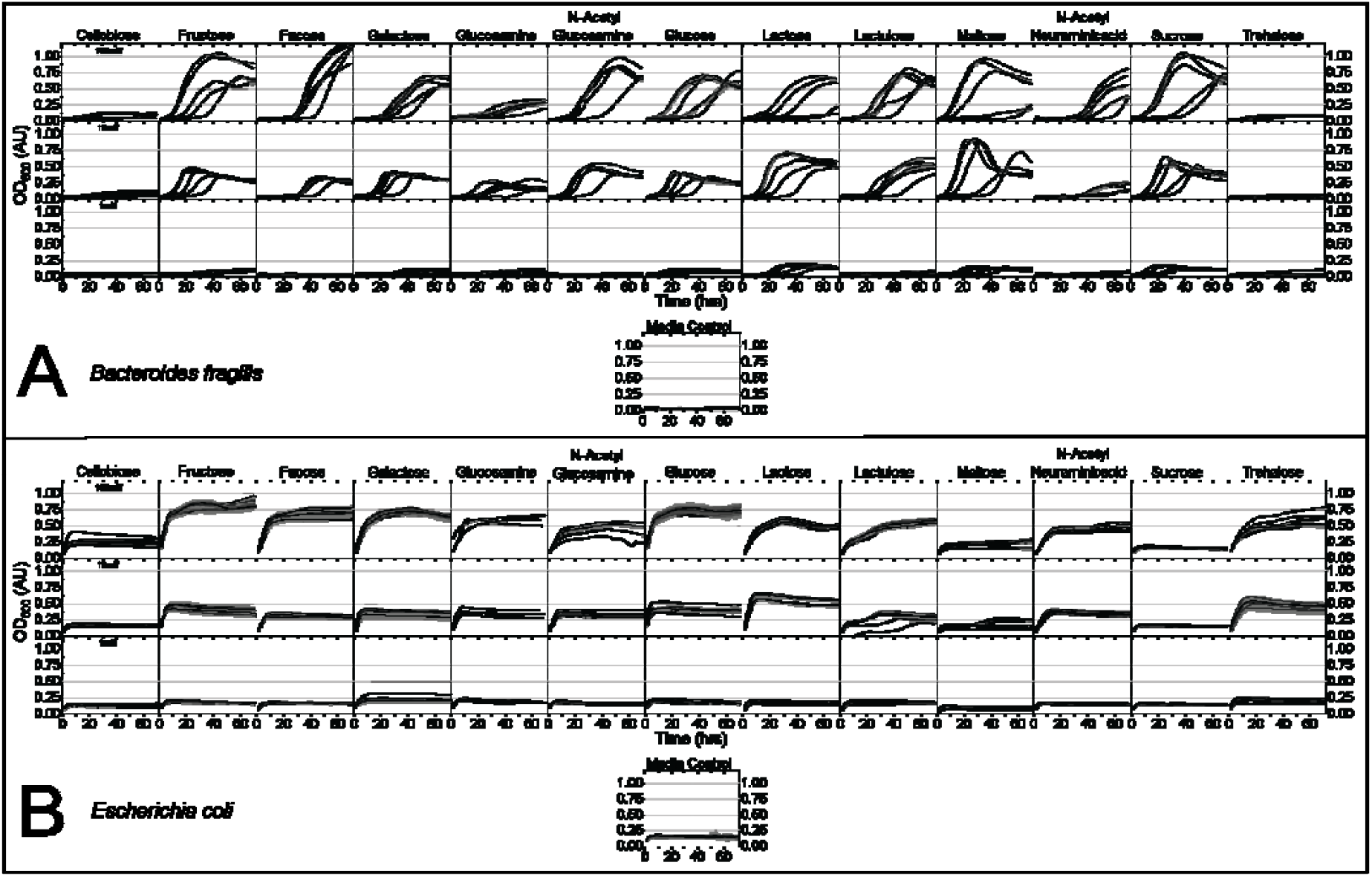
Overall growth curves of *Bacteroides fragilis* (A) and *Escherichia* coli (B) for 13 sugars at three concentrations. Each dark line represents the average OD_600_ of the replicates for an experimental run. The gray shading represents the standard deviation of the OD_600_ of the replicates. The media control was included to show the growth in the absence of a sugar.

**Figure 4.**
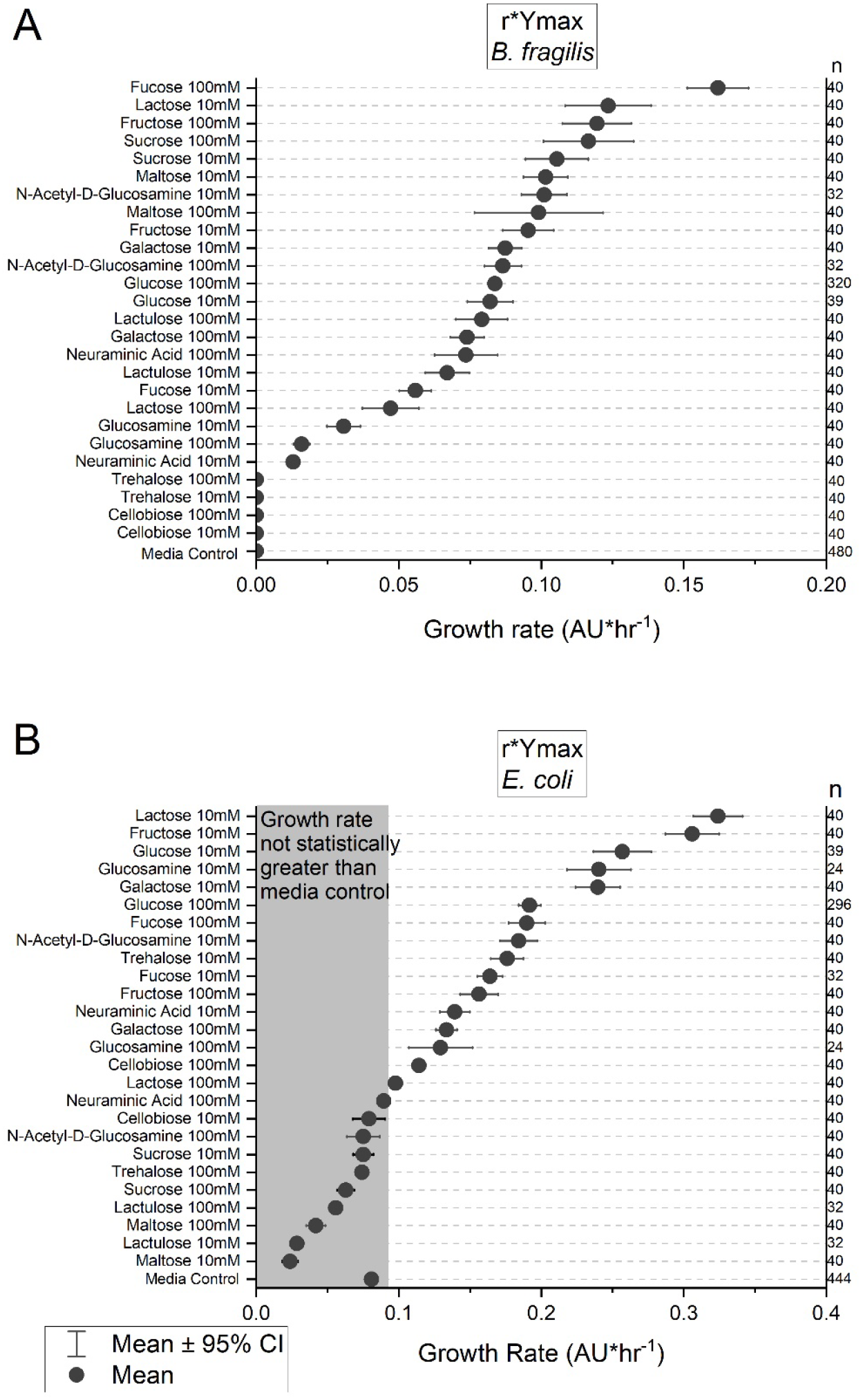
(A) Overall growth rate data of *Bacteroides fragilis*. Each dot represents the average value of a different experimental condition or the negative (media) control. The lines represent the 95% confidence interval of each condition. (B) Overall growth rate data of *Escherichia coli*. The gray shaded area represents the data that was not significantly greater than the media control. The numbers on the right are the number of replicates.

**Figure 5.**
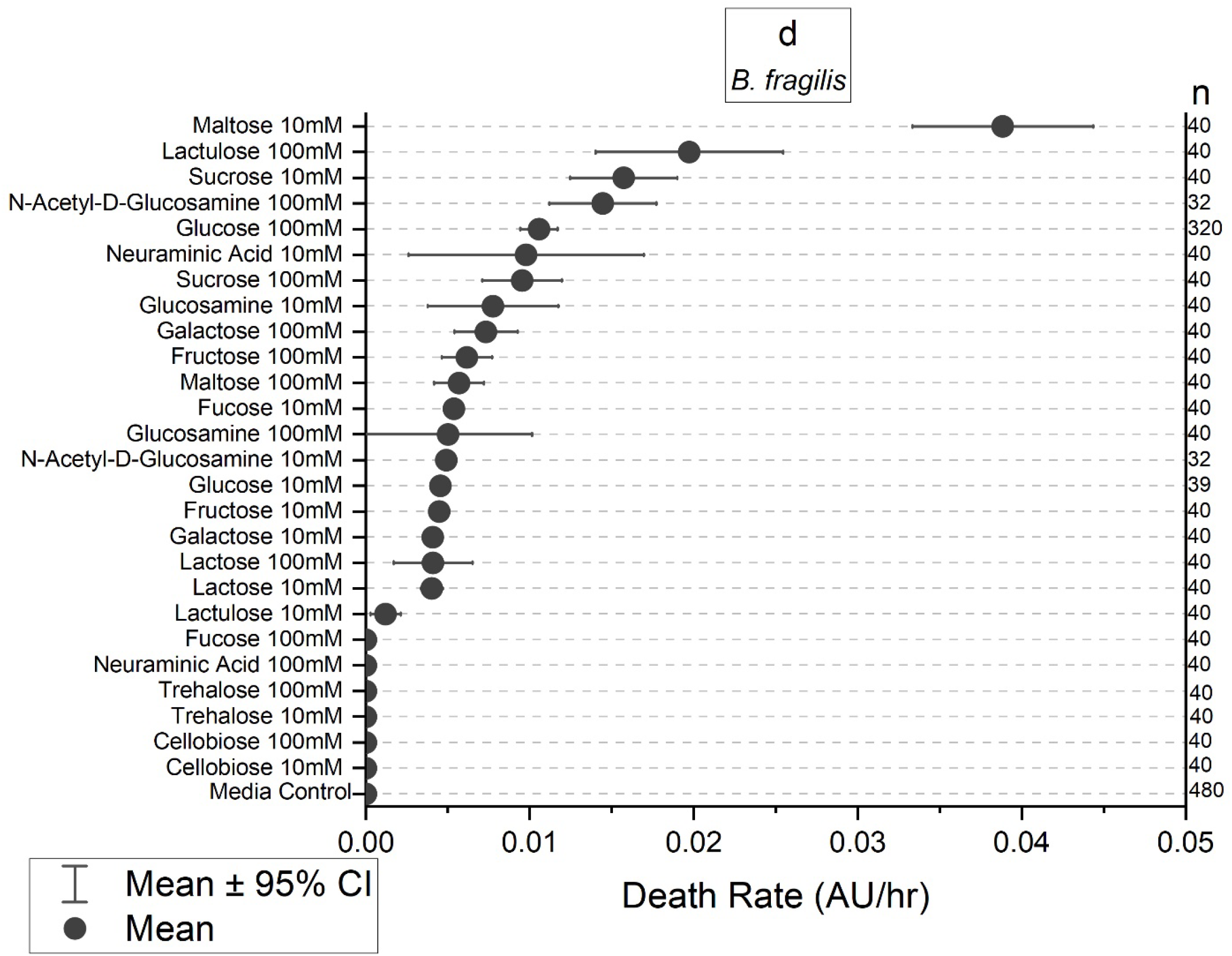
Death rate data of *Bacteroides fragilis*. Each dot represents the average value of a different experimental condition or the negative (media) control. The lines represent the 95% confidence interval of each condition. The numbers on the right are the number of replicates.

The growth was analyzed using the product of the r and Y_max_ parameter (Fig. 4). For *E. coli*, growth is led by the 10 mM concentration for most sugars except for fucose and cellobiose which are led by the 100 mM concentration. The growth of *B. fragilis* on these sugars appears to be dependent on the sugar and not the concentration. Fucose, lactose, and neuraminic acid appear to be concentration dependent as they are the only ones with large differences in their growth values between concentrations. The death rate was only looked at for *B. fragilis* in which most of the sugars produce a death effect. There is a concentration dependence for the d parameters for maltose and lactulose, which is not seen when analyzing their growth.

A similar experiment conducted by Desai *et al*. using *E. coli* showed no growth for N-acetyl neuraminic acid or cellobiose while our experiments found a statistically significant growth rate for these two sugars^30^. Similar trends as our findings are found for cellobiose, galactose, glucose, and N-acetyl glucosamine. We found growth on neuraminic acid while Desai *et al*. did not. The difference is likely because Desai *et al*. using a different strain of *E. coli* (HS). Other differences such as Desai *et al*. reporting higher growth levels for glucosamine and fucose than fructose are likely due to differences in carbohydrate usage among different strains.

The interactive web application Dashing Growth Curves was utilized for a preliminary comparison of the growth parameters of the Modified Gompertz and Modified Logistic models to our model parameters^31^. The product of r or maximum growth rate and Y_max_ was calculated for the three models. The results are not comparable due to the high standard deviation of the parameters obtained from using the web application (Table S4). The high standard deviations are likely due to the association of blanks to samples not being automated. Future work will include a more comprehensive comparison of the models.

## 4. Conclusions

This paper provides a model that can be used to predict a variety of different trends found in bacterial growth. The parameters are used to indicate a contribution to growth whether positive or negative. This model improves upon previous models in two key ways: i) it does not require knowledge of the metabolic interactions for the trends showing several growth periods, and ii) a single model allows characterization of trends in growth that are sigmoidal, non-sigmoidal, death, or a combination. A commonly used model like the Gompertz equation is said to be applicable for both death and growth but has a severe limitation of N(t=0) not equaling N(0) which may disturb curve fitting. While logistic and log-linear equations can only be applied to growth or death, respectively.

Three limitations of this model include: i) it lacks lag phase as a parameter, ii) the parameters do not indicate what is causing change in growth, and iii) r value is calculated for entire period. For example, this model is unable to determine the cause for the large death values seen in *B. fragilis* grown using maltose as the carbon source. The *B. fragilis* strain used in these experiments has prophage integrated in the genome. Experiments were conducted to induce the phage to replicate the death seen (Fig. S3). No difference was seen between the control and induced phage studies implying phage were not the source of the sudden drop in turbidity.

Investigations on chemical changes that may be causing the death are part of future studies. Calculating the r value for the entire growth period creates discrepancies when analyzing data that may have the same instantaneous growth rate, but reach Y_max_ more slowly or rapidly thus reporting a different r value. Further work includes adapting the model for several growth rates to deconvolute data that exhibits more than one growth rate before reaching Y_max_.

We anticipate that the use of models like the one developed here can provide quantification to commonly observed trends in growths. This quantification can then guide design of growth environments for microbes, for example, in bioreactors or in microbiome engineering.

## 5. Data Availability

The raw data files for this work will be available on Mendeley Data under the title “Modelling complex growth profiles of *Bacteroides fragilis* and *Escherichia coli* on various carbohydrates in an anaerobic environment.”

## 6. Acknowledgements

We are grateful to Purdue University’s Data Science Consulting Services to help with the data analysis and development of the model. We are also thankful to Ann Christine Caitlin and Steven Clark for helping us set up the analysis on DEEDS. The work was funded through Purdue’s internal funds.

## Notes

### Competing Interest Statement

MSV has interests in Krishi, Inc. Krishi, Inc. did not fund this work.

### Summary of Updates

Addition of Supporting Information.

