## Supporting Information for "Modelling complex growth profiles of *Bacteroides fragilis* and *Escherichia coli* on various carbohydrates in an anaerobic environment"

**for**

**^+^**Equal contribution

**Supporting Tables**

Table S1. Starting pH of 7.0. Concentrations are adapted from Zhang *et al.* ZMB1.(1) ^a^Necessity of each component: E = essential component (OD < 0.1); I = important component (0.1 ≤ OD < 0.4); S, somewhat important component (0.4 ≤ OD < 0.6); L = least important (or even detrimental) component (OD ≥ 0.6). These classifications were determined using the leave-one-out experiments from Zhang *et. al*. (1)

| **Variable category** | **Chemical component** | **Necessity^a^** | **Concentration** | | **Supplier** | |
| --- | --- | --- | --- | --- | --- | --- |
|  |  |  | **g/L** | **mM** | **name** | **catalog #** |
| **Essential amino acids** | L-Histidine | E | 0.17 | 1.0957 | Sigma-Aldrich | H8000 |
|  | L-Isoleucine | E | 0.24 | 1.8296 | Alfa Aesar | A13699 |
|  | L-Leucine | E | 1 | 7.6234 | Sigma-Aldrich | L8000 |
|  | L-Methionine | E | 0.06 | 0.4021 | Sigma-Aldrich | M9625 |
|  | L-Valine | E | 0.7 | 5.9753 | Sigma-Aldrich | V0500 |
|  | L-Arginine | E | 0.72 | 4.1331 | Alfa Aesar | A15738 |
| **Vitamin** | Inositol | S | 0.002 | 0.0111 | Sigma-Aldrich | 57570 |
| **Phosphate buffers** | KH_2_PO_4_ | E | 3.1 | 22.7800 | Sigma-Aldrich | P5655 |
|  | K_2_HPO_4_ | I | 6.4 | 36.7449 | Sigma-Aldrich | P8281 |
| **Other amino acid** | L-Glutamic acid | L | 0.6 | 4.0780 | Alfa Aesar | A15031 |
|  | L-Phenylalanine | L | 0.4 | 2.4214 | Alfa Aesar | A13238 |
|  | L-Proline | S | 0.7 | 6.0800 | Alfa Aesar | A10199 |
|  | L-Asparagine | S | 0.5 | 3.7845 | Sigma-Aldrich | A4159 |
|  | L-Aspartic acid | L | 0.05 | 0.3756 | Alfa Aesar | A13520 |
|  | L-Glutamine | L | 0.6 | 4.1055 | Sigma-Aldrich | G8540 |
|  | L-Serine | S | 0.5 | 4.7577 | Alfa Aesar | A11179 |
|  | L-Threonine | S | 0.5 | 4.1975 | Sigma-Aldrich | T8625 |
|  | L-Cysteine HCl | L | 0.2 | 1.2689 | Sigma-Aldrich | C1276 |
|  | L-Alanine | S | 0.4 | 4.4896 | Alfa Aesar | A15804 |
|  | Glycine | S | 0.3 | 3.9964 | Sigma-Aldrich | 50046 |
|  | L-Lysine HCl | S | 0.5 | 2.7375 | Alfa Aesar | A16249 |
|  | L-Tryptophan | L | 0.2 | 0.9793 | Sigma-Aldrich | T0254 |
|  | L-Tyrosine | S | 0.3 | 1.6557 | Alfa Aesar | A11141 |
| **Important vitamin** | Biotin | S | 0.006 | 0.0246 | Sigma-Aldrich | B4501 |
|  | Calcium pantothenate | I | 0.0012 | 0.0025 | Sigma-Aldrich | 21210 |
|  | Niacin | I | 0.0009 | 0.0073 | Sigma-Aldrich | N4126 |
|  | Pyridoxal HCl | I | 0.0048 | 0.0236 | Sigma-Aldrich | P9130 |
|  | Riboflavin | I | 0.0009 | 0.0024 | Sigma-Aldrich | R9504 |
| **Important mineral** | MgSO_4_·7H_2_O | E | 1 | 4.0574 | Alfa Aesar | 11596 |
|  | FeSO_4_·7H_2_O | S | 0.004 | 0.0144 | Alfa Aesar | A15178 |
|  | ZnSO_4_·7H_2_O | I | 0.005 | 0.0174 | Alfa Aesar | 33399 |
| **Other vitamin** | Folic acid | S | 0.00056 | 0.0013 | Sigma-Aldrich | F7876 |
|  | p-Aminobenzoic acid | S | 5.6E-05 | 0.0004 | Sigma-Aldrich | A9878 |
|  | Thiamine HCl | S | 0.00056 | 0.0017 | Sigma-Aldrich | T4625 |
| **Fatty acid** | Potassium acetate | I | 0.9 | 9.1704 | Sigma-Aldrich | 60035 |
|  | Lipoic acid | S | 0.001 | 0.0048 | Sigma-Aldrich | T5625 |
|  | Tween 80 | S | 0.5 | 0.8267 | TCI Chemicals | T0546 |
| **Nucleic acid base** | Adenine | S | 0.011 | 0.0814 | Sigma-Aldrich | A2786 |
|  | Guanine | S | 0.0056 | 0.0371 | Alfa Aesar | A12024 |
|  | Uracil | S | 0.023 | 0.2052 | Sigma-Aldrich | U0750 |
|  | Xanthine | S | 0.0038 | 0.0250 | Sigma-Aldrich | X7375 |
| **Other buffer** | MOPS | I | 15 | 71.6812 | Alfa Aesar | A12914 |
|  | Tricine | S | 1.5 | 8.3718 | Sigma-Aldrich | T0377 |
| **Trace mineral** | (NH_4_)_6_Mo_7_O_24_·4H_2_O | S | 0.00019 | 0.0002 | Alfa Aesar | A13766 |
|  | MnSO_4_·4H_2_O | S | 0.00038 | 0.0017 | Alfa Aesar | B22081 |
|  | CaCl_2_·2H_2_O | S | 0.04 | 0.2721 | Sigma-Aldrich | 21097 |
|  | CoCl_2_·6H_2_O | S | 0.00019 | 0.0008 | Sigma-Aldrich | 202185 |
|  | CuSO_4_·5H_2_O | S | 0.00019 | 0.0008 | Sigma-Aldrich | 209198 |
|  | H_3_BO_3_ | S | 0.00075 | 0.0121 | Sigma-Aldrich | 202878 |
|  | K_2_SO_4_ | S | 0.023 | 0.1320 | Alfa Aesar | 30485 |
|  | KI | S | 0.00011 | 0.0007 | Alfa Aesar | A12704 |
| **Chelator** | EDTA | S | 0.0075 | 0.0257 | Sigma-Aldrich | EDS |
|  | Nitrilotriacetic acid | S | 0.0075 | 0.0392 | Sigma-Aldrich | N9877 |
| **Other component** | Glutathione | S | 0.015 | 0.0488 | Sigma-Aldrich | G4251 |
|  | (NH_4_)_2_SO_4_ | S | 1 | 7.5681 | Sigma-Aldrich | A4418 |
|  | NaCl | S | 3 | 51.3347 | Fisher Chemical | S271 |

Table S2. Preparation of sufficient carbohydrate solutions to fill 44 wells using the procedure detailed in this manuscript (sufficient for 4 experiments).

| **Preparation of 11 mL of 111 mM carbohydrate solutions in ZMB1** | | | |
| --- | --- | --- | --- |
| **Chemical component** | **Amount weighed (g)** | **Supplier** | |
|  |  | **Name** | **catalog #** |
| D-Glucose | 0.220 | Sigma-Aldrich | G8270 |
| D-Galactose | 0.220 | Sigma-Aldrich | G0625 |
| D-Fructose | 0.220 | Sigma-Aldrich | F3510 |
| L-Fucose | 0.200 | Fisher Scientific | AC225880250 |
| N-Acetyl-D-Glucosamine | 0.270 | Sigma-Aldrich | A3286 |
| D-Glucosamine HCl | 0.268 | Fisher Scientific | AAA1553218 |
| N-Acetylneuraminic Acid | 0.378 | Activate Scientific | AS23626 |
| D-Maltose·H_2_O | 0.440 | Alfa Aesar | A16266 |
| D-Lactose·H_2_O | 0.440 | Sigma-Aldrich | L254 |
| D-Cellobiose | 0.418 | Sigma-Aldrich | 22150 |
| D-Sucrose | 0.418 | Alfa Aesar | A15583 |
| D-Lactulose | 0.418 | Alfa Aesar | J60160 |
| D-Trehalose·2H_2_O | 0.462 | Alfa Aesar | A19434 |

Table S3. Preliminary comparison of the growth rate (the product of r or µ_max_ and Y_max_) from our model to the modified Logistic and modified Gompertz models using the web application Dashing Growth Curves(2).

| **Sugar and Concentration (mM)** | **Growth Rate (AU*hr⁻¹)** | | |
| --- | --- | --- | --- |
|  | **Ours** | **Logistic** | **Gompertz** |
| Lactose - 100 | 0.098 ± 0.0151 | 0.242 ± 0.04 | 0.277 ± 0.04 |
| Cellobiose-100 | 0.114 ± 0.0148 | 0.3 ± 0.156 | 0.216 ± 0.098 |
| Trehalose - 10 | 0.176 ± 0.0356 | 0.868 ± 0.22 | 1.093 ± 0.341 |
| Glucose - 100 | 0.192 ± 0.0677 | 0.569 ± 0.0354 | 0.642 ± 0.042 |
| Fructose - 10 | 0.306 ± 0.0586 | 1.67 ± 1.318 | 0.377 ± 0.038 |
| Lactose -10 | 0.324 ± 0.0542 | 1.437 ± 0.309 | 1.29 ± 0.286 |

**Supporting Figures**

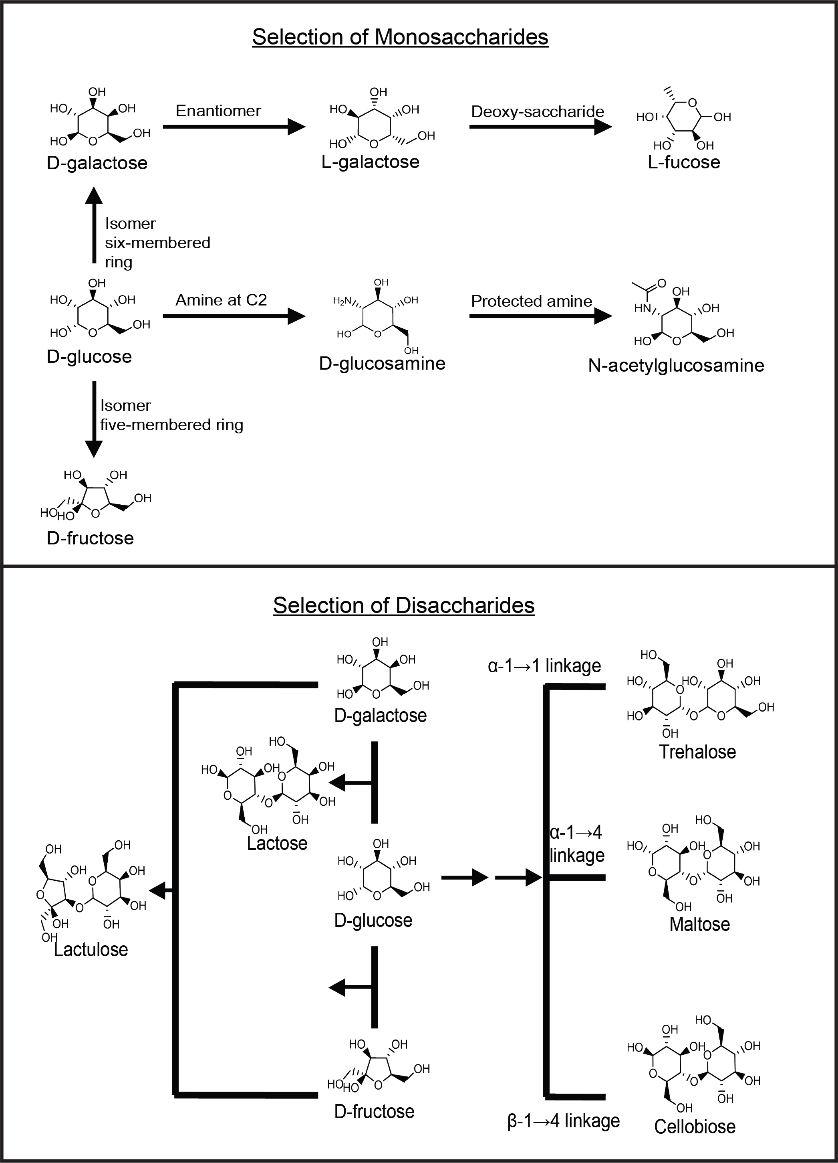

Figure S1. Schematic of the reasoning for the selection of the 13 sugar sources. Selection was based around glucose and focused on its isomers, the removal or introduction of functional groups, the different glycosidic linkages, and utilizing two different monosaccharides.

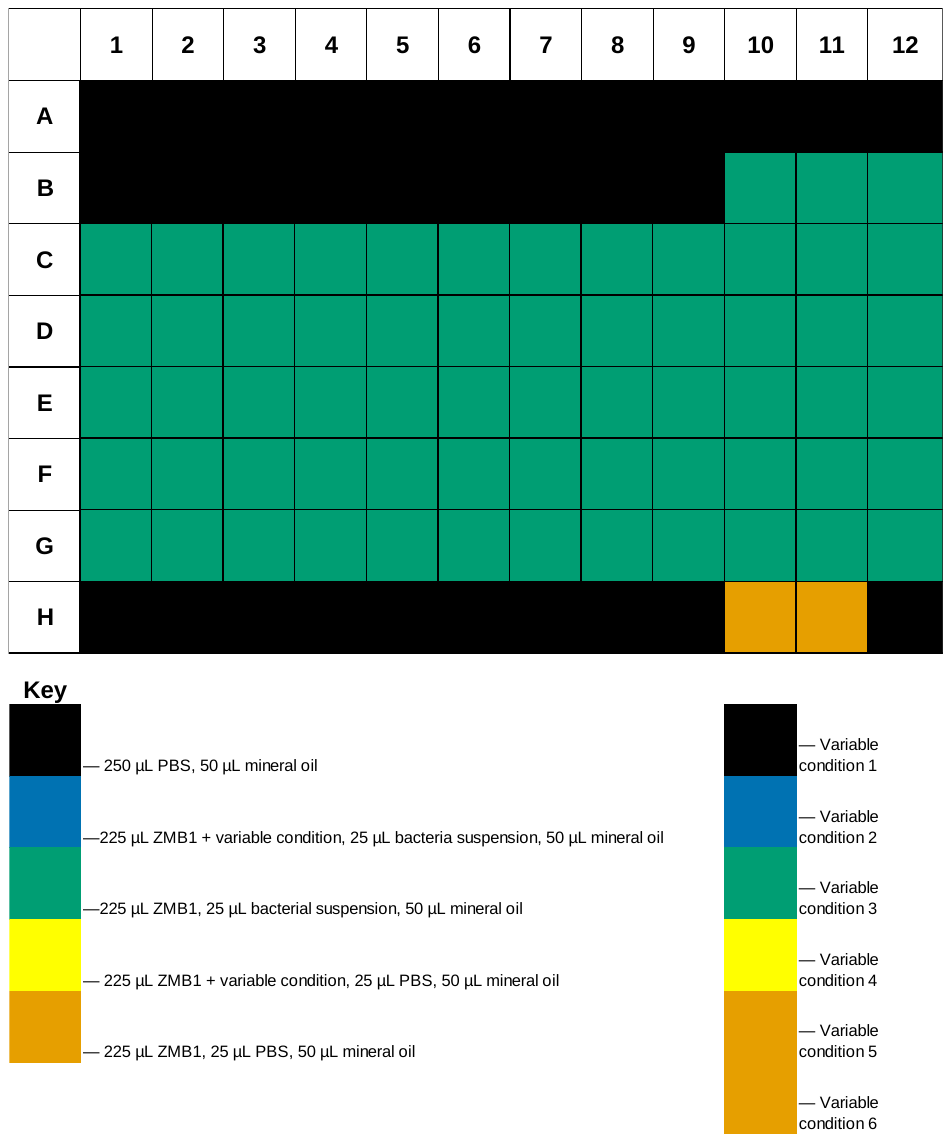

Figure S2. Experimental setup using a 72-wells in a 96-well plate. Each row B-F represent a different experimental condition. Columns 1 and 12 serve as the blank for their row's respective experimental condition. Columns 10 and 11 serve as the negative control. Wells H10-11 serve as the blank for the negative control. Variable condition refers to the specific concentration of a sugar. Variable condition 6 was set to glucose 111mM for each plate as a positive control.

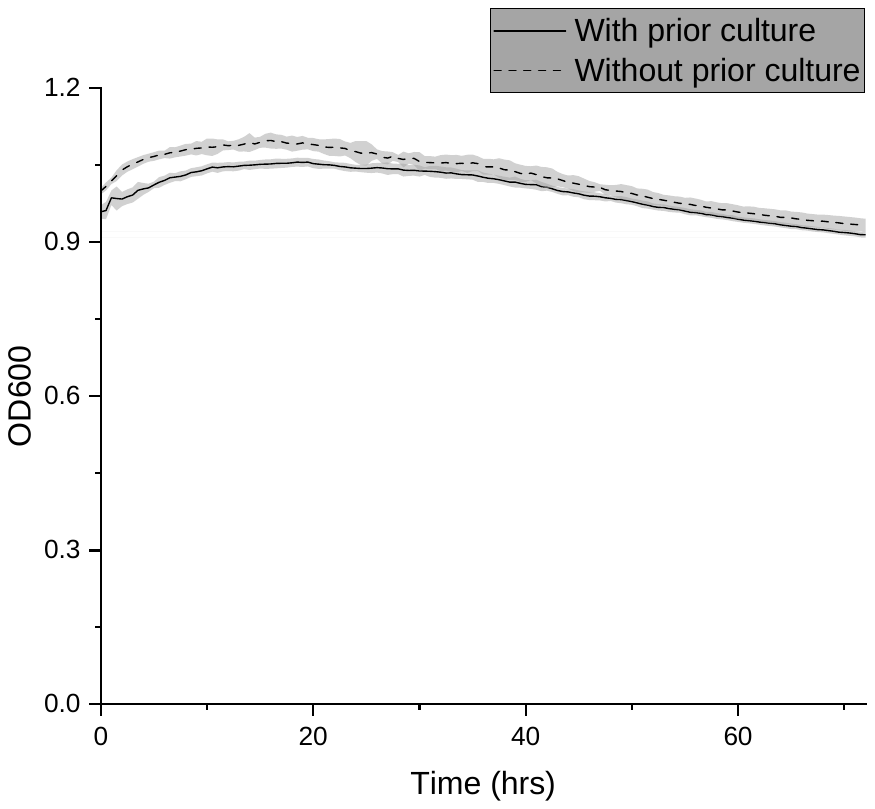

Figure S3. Death of *B. fragilis* grown in ZMB1 with and without the addition of a sample of prior 10 mM maltose culture during death phase to test for the presence of a phage. Each culture was grown to an OD of ~1. In a 96-well plate, 25 µl of the 10mM maltose culture at the death stage was added. The sample without the prior culture had 25 µl of PBS.

**Supporting Scripts**

DEEDS parsing script: parsing.py

\# -*- coding: utf-8 -*-

"""

Created on Sun Nov 25 16:12:41 2018

@author: Moiz

"""

import os

import sys

import pandas as pd

import numpy as np

import matplotlib.pyplot as plt

import math

import colorsys

import datetime

import argparse

import collections

import re

import shutil

import copy

posContrName = "Contr+"

negContrName = "Contr-"

CONTROL_KEY = "controls"

DATA_KEY = "data"

global data

global time

global nameToCell

global codebook

global colorBook

global uselegend

def init_args():

parser = argparse.ArgumentParser(description="vermalab raw data parser and visualizer")

parser.add_argument("rawdatafile", help="raw_data.xlsx file; if you're having issues, try not having a space in the file name")

parser.add_argument("labelsfile", help="labels.xlsx file; same troubleshooting tip as above")

return parser.parse_args()

def main():

args = init_args()

if not os.path.exists(args.rawdatafile):

raise ValueError("raw data file does not exist")

if not os.path.exists(args.labelsfile):

raise ValueError("labels file does not exist")

exp = Experiment(args.rawdatafile, args.labelsfile)

exp.save_all_combinations()

exp.explodeData()

class Experiment:

def __init__(self, rawdatafile, labelsfile, colorsfile=None):

if not os.path.exists(rawdatafile):

raise ValueError("raw data file does not exist")

if not os.path.exists(labelsfile):

raise ValueError("labels file does not exist")

if colorsfile:

self.add_colors_file(colorsfile)

dataInfo = os.path.splitext(os.path.basename(rawdatafile))[0].split('_')

labelInfo = os.path.splitext(os.path.basename(labelsfile))[0].split('_')

if (labelInfo[0] == dataInfo[0]) and (labelInfo[1] == dataInfo[1]):

self.experimentDate = labelInfo[0]

self.bacteria	= labelInfo[1]

else:

raise ValueError("labels file and raw data file names do not match")

self.data, self.time = load_raw_data(rawdatafile)

self.labels = load_labels(labelsfile)

self.colorbook = dict()

def add_colors_file(self, colorsfile):

if not os.path.exists(colorsfile):

raise ValueError("colors file does not exist")

\# self.colors =

def get_one_run_raw(self, freq, compound, concentration):

if not isinstance(freq, str):

raise TypeError("frequency must be a string")

if not isinstance(compound, str):

raise TypeError("compound must be a string type")

if not isinstance(concentration, str):

raise TypeError("concentration must be a string type")

data = self.data[freq][self.labels[compound][concentration][DATA_KEY]]

control = self.data[freq][self.labels[compound][concentration][CONTROL_KEY]]

time = self.time[freq]

return time, data, control

def get_one_run(self, freq, compound, concentration):

time, data, control = self.get_one_run_raw(freq, compound, concentration)

subbed = subControls(control, data)

avgs, stds = avgDataAndStd(subbed)

time = fixTime(time)

name = freq + "_" + compound + "_" + concentration

return pd.Series(time).to_frame(name + "_time"), avgs.to_frame(name), stds.to_frame(name + "_err")

def collect_runs(self, frequencies, compounds, concentrations):

if len(frequencies) != len(compounds) or len(frequencies) != len(concentrations):

raise ValueError("Inputs must be iterables of the same length")

minlen = min([self.data[freq].shape[0] for freq in frequencies])

times = pd.DataFrame()

avgs = pd.DataFrame()

stds = pd.DataFrame()

def addToDf(dft, df):

if dft.empty: return df

else: return dft.join(df)

for i in range(len(frequencies)):

time, avg, std = self.get_one_run(frequencies[i], compounds[i], concentrations[i])

times = addToDf(times, time.loc[0:minlen])

avgs = addToDf(avgs, avg.loc[0:minlen])

stds = addToDf(stds, std.loc[0:minlen])

return times, avgs, stds

def plot(self, ax, times, avgs, stds):

if times.shape[1] != avgs.shape[1] or times.shape[1] != stds.shape[1]:

raise ValueError("times, avgs, and stds must have the same number of columns")

for i in range(0, len(avgs.columns)):

avg = avgs[avgs.columns[i]]

std = stds[stds.columns[i]]

time = times[times.columns[i]]

freq, comp, conc = split_name(avgs.columns[i])

hsv = color_mapper(self.colorbook, comp)

rgb = adjust_color(hsv, int(conc))

time = time.to_numpy()

avg = avg.to_numpy()

std = std.to_numpy()

if time.shape[0] != avg.shape[0]:

print("WARNING: time and avg in " + avgs.columns[i] + " are not same length", file=sys.stderr)

if std.shape[0] != avg.shape[0]:

print("WARNING: std and avg in " + avgs.columns[i] + " are not same length", file=sys.stderr)

minlen = min([var.shape[0] for var in [time, std, avg]])

time = time[:minlen]

std = std[:minlen]

avg = avg[:minlen]

ax.plot(time, avg, label=avgs.columns[i], color=rgb)

ax.fill_between(time, avg - std / 2, avg + std / 2, color=(rgb[0], rgb[1], rgb[2], .5))

\# chartBox = ax.get_position()

\# ax.set_position([chartBox.x0, chartBox.y0, chartBox.width*0.75, chartBox.height])

\# ax.legend(loc="upper center", bbox_to_anchor=(1.45, 0.8))

ax.legend()

def graph(self, title, times, avgs, stds, xlabel="Time (hrs)", ylabel="OD"):

fig = plt.figure()

ax = fig.gca()

self.plot(ax, times, avgs, stds)

ax.set_title(title)

ax.set_ylabel(ylabel)

ax.set_xlabel(xlabel)

return fig

def save_csv(self, name, times, avgs, stds):

data = times.join(avgs).join(stds)

data.to_csv(name + ".csv")

def save_fig(self, title, fig):

fig.savefig(title + ".png")

def save_all_combinations(self):

for freq in self.data.keys():

if os.path.exists(freq):

shutil.rmtree(freq)

os.mkdir(freq)

for comp in self.labels:

if comp == "ZMB1":

continue

os.mkdir(os.path.join(freq, comp))

concs = list(self.labels[comp].keys())

comps = [comp] * len(concs) + ["ZMB1"]

concs += ["1"]

freqs = [freq] * len(concs)

times, avgs, stds = self.collect_runs(freqs, comps, concs)

title = os.path.join(freq, comp, freq + "_" + comp + "_concs")

print(title)

fig = self.graph("test", times, avgs, stds)

self.save_fig(title, fig)

self.save_csv(title, times, avgs, stds)

plt.close(fig)

def explodeData(self):

plateRowColumnPattern = "^(\d+)([A-Z]+)(\d+)"

controls = {}

for waveLength in self.data:

controls[waveLength] = {}

for carbonSource in self.labels:

controls[waveLength][carbonSource] = {}

for concentration in self.labels[carbonSource]:

controls[waveLength][carbonSource][concentration] = {}

nTime	= -1

nData	= -1

explodedHeader = "%s,%s,%s,%s,%s" % ('Date','Bacteria',

'Wavelength','Carbon Source','Concentration')

plateWells = self.labels[carbonSource][concentration]['data'] + \

self.labels[carbonSource][concentration]['controls']

natsort(plateWells)

for plateWell in plateWells:

plate,wellRow,wellColumn = re.search(plateRowColumnPattern,plateWell).groups()

if plateWell in self.labels[carbonSource][concentration]['controls']:

if not plate in controls[waveLength][carbonSource][concentration]:

controls[waveLength][carbonSource][concentration][plate] = []

controls[waveLength][carbonSource][concentration][plate].append(self.data[waveLength][plateWell])

explodedHeader += ",%s,%s,%s,%s,%s,%s,%s,%s" % ('Replicate', \

'Plate','Well Row','Well Column','Bacteria Present', \

'Time','Value','Adjusted Value')

if nTime < 0:

nTime = len(self.time[waveLength][plate])

nData = len(self.data[waveLength][plateWell])

else:

nTime = max(nTime,len(self.time[waveLength][plate]))

nData = max(nData,len(self.data[waveLength][plateWell]))

for plate in controls[waveLength][carbonSource][concentration]:

nSamples = len(controls[waveLength][carbonSource][concentration][plate])

\#	print("%s %s %s PLATE = %s, %d" % (waveLength,carbonSource,concentration,plate,nSamples))

sample = 0

averages = copy.copy(controls[waveLength][carbonSource][concentration][plate][sample])

sample += 1

while sample < nSamples:

averages = [total+value for total,value in zip(averages,controls[waveLength][carbonSource][concentration][plate][sample])]

sample += 1

controls[waveLength][carbonSource][concentration][plate] = [total/nSamples for total in averages]

explodedFile = "%s_%s_%s_%s_%s.csv" % (self.experimentDate,self.bacteria,

waveLength,carbonSource,concentration)

with open(explodedFile,'w') as fp:

fp.write(explodedHeader + '\n')

for dataIndex in range(nTime):

\#	averageIndex = dataIndex

averageIndex = 0

explodedRow = "%s,%s,%s,%s,%s" % (self.experimentDate,self.bacteria,

waveLength,carbonSource,concentration)

replicate = 0

for plateWell in plateWells:

if plateWell in self.labels[carbonSource][concentration]['data']:

replicate += 1

plate,wellRow,wellColumn = re.search(plateRowColumnPattern,plateWell).groups()

bacteriaPresent = 1

try:

dataTime = self.time[waveLength][plate][dataIndex]

dataValue = self.data[waveLength][plateWell][dataIndex]

averageControlValue = controls[waveLength][carbonSource][concentration][plate][averageIndex]

if dataValue >= 0.:

adjustedValue = dataValue - averageControlValue

else:

adjustedValue = dataValue

except:

explodedRow += ",%d,%s,%s,%s,%d,,," % (replicate, \

plate,wellRow,wellColumn,bacteriaPresent)

else:

explodedRow += ",%d,%s,%s,%s,%d,%f,%f,%f" % (replicate, \

plate,wellRow,wellColumn,bacteriaPresent,

dataTime,dataValue,adjustedValue)

else:

plate,wellRow,wellColumn = re.search(plateRowColumnPattern,plateWell).groups()

bacteriaPresent = 0

try:

dataTime = self.time[waveLength][plate][dataIndex]

dataValue = self.data[waveLength][plateWell][dataIndex]

averageControlValue = controls[waveLength][carbonSource][concentration][plate][averageIndex]

if dataValue >= 0.:

adjustedValue = dataValue - averageControlValue

else:

adjustedValue = dataValue

except:

explodedRow += ",NA,%s,%s,%s,%d,,," % (plate,wellRow,wellColumn,bacteriaPresent)

else:

explodedRow += ",NA,%s,%s,%s,%d,%f,%f,%f" % (plate,wellRow,wellColumn,bacteriaPresent,

dataTime,dataValue,adjustedValue)

fp.write(explodedRow + '\n')

def natsort(listValues):

""" Sort the given list in the way that humans expect.

"""

def tryint(stringValue):

try:

return int(stringValue)

except:

return stringValue.lower()

def alphanum_key(stringValue):

""" Turn a string into a list of string and number chunks.

"z23a" -> ["z", 23, "a"]

"""

return [ tryint(c) for c in re.split('([0-9]+)', stringValue) ]

listValues.sort(key=alphanum_key)

def split_name(name):

pattern = "^(\d+)_([\w\-]+)_(\d+)"

res = re.search(pattern, name)

return res[1], res[2], res[3]

def adjust_color(hsv, concentration):

percent = math.log10(float(concentration) * 10)

h, s, v = hsv

v = ((1 - .3) / 3) * percent + .3

if v > 1:

v = 1

r, g, b = colorsys.hsv_to_rgb(h, s, v)

return (r,g,b)

def color_mapper(colorbook, compound):

if compound in colorbook:

return colorbook[compound]

else:

colorbook[compound] = np.random.rand(1, 3)[0]

return colorbook[compound]

def load_labels(filename):

labels = pd.ExcelFile(filename)

def defaultdefaultcallable():

return {CONTROL_KEY: list(), DATA_KEY: list()}

def defaultcallable():

return collections.defaultdict(defaultdefaultcallable)

codebook = collections.defaultdict(defaultcallable)

pattern = "([\w\-]+)_(\d+)_(\d)"

for x in labels.sheet_names:

df = labels.parse(x)

plateno = x.split(' ')[1]

def startcond(dataframe, i):

return dataframe.iloc[i, 1] == 1 and dataframe.iloc[i, 2] == 2

def endcond(dataframe, i):

return isinstance(dataframe.iloc[i, 1], float) and math.isnan(dataframe.iloc[i, 1])

colrange = [1, 13]

dataframe = getDFOI(df, 0, df.shape[0], startcond, endcond, colrange)

for row in dataframe.index:

for col in dataframe.columns:

\# print(row, col)

if isinstance(dataframe[col].loc[row], str):

res = re.search(pattern, dataframe[col].loc[row])

key = plateno + chr(ord("A") + row) + str(col)

if res[3] == "1":

codebook[res[1]][res[2]][DATA_KEY].append(key)

else:

codebook[res[1]][res[2]][CONTROL_KEY].append(key)

return codebook

def load_raw_data(filename):

data = pd.ExcelFile(filename)

\# totalData = pd.DataFrame()

totaldata = collections.defaultdict(pd.DataFrame)

\# totalTime = pd.DataFrame()

totaltime = collections.defaultdict(pd.DataFrame)

for x in data.sheet_names:

df = data.parse(x)

plateno = x.split(' ')[1]

if not plateno.isdigit():

raise ValueError('Sheet Names must be of the following form: "Plate <number> ..."\n'

'important that integer value is placed after "Plate"')

\# array of indexes at which the frequencies are listed in the first column

freq_indexes = df[df[df.columns[0]].apply(lambda x: isinstance(x, int))].index

\# uses freq_indexes to extract the table of each frequency

def startcond(dataframe, i):

return dataframe.iloc[i][1] == 'Time' and dataframe.iloc[i][3] == 'A1'

def endcond(dataframe, i):

if not (isinstance(dataframe.iloc[i][1], datetime.datetime) or isinstance(dataframe.iloc[i][1], datetime.time)):

return True

if i > 1 and datetimeToHours(dataframe.iloc[i][1]) == 0:

return True

colrange = [1, 99]

dfs = {str(df.iloc[freqind][0]):

getDFOI(df, freqind, df.shape[0], startcond, endcond, colrange, plateno)

for freqind in freq_indexes}

\# df = getDFOI(df, plateno)

times = {freq: convertDateTimeSeries(val[val.columns[0]], plateno) for freq, val in dfs.items()}

\# time = convertDateTimeSeries(df[df.columns[0]], plateno)

rawdatas = {freq: val[val.columns[2:]].replace("OVRFLW", -1) for freq, val in dfs.items()}

\# rawdata = df[df.columns[2:]]

for f in dfs:

if totaldata[f].empty:

totaldata[f] = rawdatas[f]

totaltime[f] = times[f]

else:

minlength = min(totaltime[f].shape[0], times[f].shape[0])

totaldata[f] = totaldata[f].iloc[0:minlength, :].join(rawdatas[f].iloc[0:minlength, :])

totaltime[f] = totaltime[f].iloc[0:minlength, :].join(times[f].iloc[0:minlength, :])

return totaldata, totaltime

def getDFOI(dataframe, startrow, maxrow, startcond, endcond, colrange, platenum: str=None):

startind = 0

endind = 0

for i in range(startrow, maxrow):

\# print(dataframe.iloc[i, :])

if startcond(dataframe, i):

startind = i

break

for j in range(startind + 1, dataframe.shape[0]):

if endcond(dataframe, j):

endind = j

break

newcolnames = dataframe.iloc[startind, 1:].to_numpy()

if platenum:

newcolnames = [platenum + val for val in newcolnames]

oldcolnames = dataframe.columns[1:]

colnamesdict = {old: new for old, new in zip(oldcolnames, newcolnames)}

indexnamesdict = {old: new for old, new in zip(np.arange(startind + 1, endind), np.arange(0, endind - startind))}

dfoi = dataframe.iloc[(startind+1):endind, colrange[0]:colrange[1]]

return dfoi.rename(columns=colnamesdict, index=indexnamesdict)

'''combine values from collectCompConc into one dataframe'''

def combineCompConc(time, avgs, stds):

df = pd.DataFrame(time, columns=["time"]).join(avgs).join(stds)

return df

'''takes data frame data and calulates mean and standard deviation across rows and returns the two series'''

def avgDataAndStd(data):

avgData = data.mean(axis=1)

stdData = data.std(axis=1)

return avgData, stdData

'''takes the dataframe of control cells, averages them, and subtracts the values row wise from data'''

def subControls(controls, data):

\# print("Subtracting Controls...")

avgControls = controls.mean(axis = 1)

avgControls = [avgControls[0]] * len(avgControls)

return data.sub(avgControls, axis=0)

def fixTime(allTime):

"""

takes all the time columns from the different plates and averages them

:param allTime: must a dataframe

:return: a numpy array

"""

\# print("Fixing Time...")

"""

avgDiffs = 0

avgStart = 0

for col in allTime:

time = allTime[col]

avgStart += time[0]

for i in np.arange(0, time.size - 1):

avgDiffs += time[i + 1] - time[i]

avgDiffs /= (allTime.shape[0] - 1) * allTime.shape[1]

avgStart /= allTime.shape[1]

"""

timenum = allTime.to_numpy(np.float64)

avgDiffs = np.average(timenum[1:] - timenum[:-1])

avgStart = np.average(timenum[0])

topVal = np.ceil(avgStart + avgDiffs * timenum.shape[0])

return np.arange(avgStart, topVal, avgDiffs)[:timenum.shape[0]]

def convertDateTimeSeries(ser, plate:str):

for i in np.arange(0, ser.shape[0]):

ser[i] = datetimeToHours(ser[i])

return ser.to_frame(plate)

def datetimeToHours(val):

dayhours = 0

if isinstance(val, datetime.datetime):

dayhours = val.day * 24

return dayhours + val.hour + (val.minute / 60) + (val.second / 3600)

'''create color book from file'''

def createcolorbook(colorFile):

global colorBook

colorBook = {}

if os.path.exists(colorFile):

colorFile = pd.ExcelFile(colorFile)

if len(colorFile.sheet_names) > 1:

print("Program only accepts label workbooks with one sheet at the moment")

return

df = colorFile.parse(colorFile.sheet_names[0])

df.columns = df.columns.str.lower()

for row, val in df.iterrows():

compName = df["compound"][row]

r = float(df["r"][row]) / 255

g = float(df["g"][row]) / 255

b = float(df["b"][row]) / 255

colorBook[compName] = colorsys.rgb_to_hsv(r, g, b)

if not colorBook[posContrName]:

colorBook[posContrName] = (1, 1, 1)

if not colorBook[negContrName]:

colorBook[negContrName] = (.5, .5, .5)

print("sucess")

return colorBook

if __name__ == '__main__':

\# data, time = load_raw_data("20191205_BF001EC001_Spectra.xlsx")

\# labels = load_labels("20191205_BF001EC001_Spectra_Labels.xlsx")

plt.ioff()

main()

DEEDS model analysis script: modelFit.py

import sys

import os

import re

import argparse

import pandas

import numpy as np

import matplotlib

matplotlib.use("Agg")

import matplotlib.pyplot as plt

from scipy import optimize

from collections import deque

import math

def fitting(replicate,

t,

y,

resetDonTmax,

rMinimum,

rMaximum,

dMinimum,

dMaximum,

iterMaximum):

ind = np.argmax(y)

tmax = t[ind]

ymax = y[ind]

dt = t[1] - t[0]

dy = np.gradient(y,dt)

message = 'Exception occurred'

objFunction = -1.

def loss(para):

r,d = para

dy_appr = r*y*(1-y/(ymax-d*(t-tmax)))

return np.mean((dy - dy_appr)**2)

bound = [(rMinimum,rMaximum),(dMinimum,dMaximum)]

try:

result = optimize.dual_annealing(loss,bound,maxiter=iterMaximum)

except:

r	= None

d	= None

tail_slope_std = None

tlag	= None

else:

message	= result.message[0]

objFunction = result.fun

r,d	= result.x

if resetDonTmax:

if ind+1 == t.size:

d = 0.

if ind < t.size-20:

tail_slope_std = np.std((y[ind+10:]-ymax)/(t[ind+10:]-tmax))

else:

tail_slope_std = 0.

with np.errstate(divide='raise'):

try:

tlag = tmax - 4./r

except FloatingPointError:

tlag = 0.

dy_appr = r*y*(1-y/(ymax-d*(t-tmax)))

y_appr = np.append([y[0]],y[:-1] + dy_appr[:-1]*dt)

plt.plot(t,y,'k.',markersize=2)

plt.plot(t,y_appr,label=replicate)

return r,d,tmax,ymax,tail_slope_std,tlag,objFunction,message

def fitModelToExperiment(experimentResultFile,

timeCutoff,

resetDonTmax,

averageHalfWidth,

rMinimum,

rMaximum,

dMinimum,

dMaximum,

iterMaximum):

results = deque()

rawDataFrame = pandas.read_csv(experimentResultFile)

columnNames = rawDataFrame.columns.to_list()

date	= rawDataFrame['Date'][0]

bacteria	= rawDataFrame['Bacteria'][0]

wavelength	= rawDataFrame['Wavelength'][0]

carbonSource = rawDataFrame['Carbon Source'][0]

concentration = rawDataFrame['Concentration'][0]

firstReplicate	= columnNames.index('Replicate')

firstPlate	= columnNames.index('Plate')

firstWellRow	= columnNames.index('Well Row')

firstWellColumn	= columnNames.index('Well Column')

firstBacteriaPresent = columnNames.index('Bacteria Present')

firstTime	= columnNames.index('Time')

firstValue	= columnNames.index('Adjusted Value')

secondReplicate	= columnNames.index('Replicate.1')

columnsPerReplicate = secondReplicate-firstReplicate

nReplicates	= (len(columnNames)-firstReplicate)//columnsPerReplicate

\# print(firstReplicate)

\# print(secondReplicate)

\# print(columnsPerReplicate)

\# print(nReplicates)

reDateTime = re.compile("(.*)(_[0-9]{8}-[0-9]{6})$")

experiment,_ = os.path.splitext(os.path.basename(experimentResultFile))

dateTimeMatch = reDateTime.match(experiment)

while dateTimeMatch:

experiment = dateTimeMatch.group(1)

dateTimeMatch = reDateTime.match(experiment)

replicateDataFrame = pandas.DataFrame()

fig = plt.figure()

nResults = 0

for k1 in range(0,nReplicates):

replicate	= rawDataFrame.iloc[0,firstReplicate+k1*columnsPerReplicate]

plate	= rawDataFrame.iloc[0,firstPlate+k1*columnsPerReplicate]

wellRow	= rawDataFrame.iloc[0,firstWellRow+k1*columnsPerReplicate]

wellColumn	= rawDataFrame.iloc[0,firstWellColumn+k1*columnsPerReplicate]

bacteriaPresent = rawDataFrame.iloc[0,firstBacteriaPresent+k1*columnsPerReplicate]

\#	print(date,bacteria,wavelength,carbonSource,concentration,replicate,plate,wellRow,wellColumn,bacteriaPresent)

if bacteriaPresent == 1:

t = np.array(rawDataFrame.iloc[:,firstTime+k1*columnsPerReplicate])

y = np.array(rawDataFrame.iloc[:,firstValue+k1*columnsPerReplicate])

if timeCutoff > 0.:

t = t[t <= timeCutoff]

y = y[:t.shape[0]]

if averageHalfWidth > 0:

windowSize = 2*averageHalfWidth+1

window = np.ones(int(windowSize))/float(windowSize)

y = np.convolve(y,window,mode='valid')

t = t[averageHalfWidth:]

t = t[:y.shape[0]]

if np.unique(y).shape[0] > 1:

nColumns = replicateDataFrame.shape[1]

if nColumns == 0:

replicateDataFrame.insert(nColumns,"Time",t,False)

nColumns += 1

replicateDataFrame.insert(nColumns,"Replicate %d" % (k1),y,False)

r,d,tmax,ymax,tail_slope_std,tlag,objFunction,fitMessage = fitting(replicate,t,y,

resetDonTmax,

rMinimum,rMaximum,

dMinimum,dMaximum,

iterMaximum)

if r != None:

nResults += 1

rymax = r*ymax

results.append([date,bacteria,wavelength,carbonSource,concentration] +

[replicate,plate,wellRow,wellColumn,bacteriaPresent,r,d,tmax,ymax,tail_slope_std,tlag,rymax,objFunction,fitMessage])

print("	%d %d %s %d : %f %f %f %f %f %f %f %f %s" % (replicate,plate,wellRow,wellColumn, \

r,d,tmax,ymax,tail_slope_std,tlag, \

rymax,objFunction,fitMessage))

else:

rymax = -1

results.append([date,bacteria,wavelength,carbonSource,concentration] +

[replicate,plate,wellRow,wellColumn,bacteriaPresent,-1,-1,tmax,ymax,-1,-1,rymax,objFunction,fitMessage])

print("	%d %d %s %d : %f %f %f %f %f %f %f %f %s" % (replicate,plate,wellRow,wellColumn, \

-1,-1,tmax,ymax,-1,-1, \

rymax,objFunction,fitMessage))

else:

ind = np.argmax(y)

tmax = t[ind]

ymax = y[ind]

rymax = -1

objFunction = -1.

fitMessage = 'Colinear data'

results.append([date,bacteria,wavelength,carbonSource,concentration] +

[replicate,plate,wellRow,wellColumn,bacteriaPresent,-1,-1,tmax,ymax,-1,-1,rymax,objFunction,fitMessage])

print("	%d %d %s %d : %f %f %f %f %f %f %f %f %s" % (replicate,plate,wellRow,wellColumn, \

-1,-1,tmax,ymax,-1,-1, \

rymax,objFunction,fitMessage))

if nResults > 0:

columns = replicateDataFrame.columns[1:]

replicateDataFrame['Average'] = replicateDataFrame[columns].mean(axis=1)

replicateDataFrame.to_csv(experiment + '_averages.csv',index=False)

plt.xlabel('time')

plt.ylabel('value')

plt.xlim(left=0)

plt.ylim(bottom=0)

plt.legend(loc='center left',bbox_to_anchor=(1, 0.5),ncol=math.ceil(nResults/20.))

plt.savefig(experiment + '.png',bbox_inches='tight')

plt.close()

return results

if __name__ == '__main__':

parser = argparse.ArgumentParser()

parser.add_argument("--timeCutoff",type=float,default=0.,required=False)

parser.add_argument("--rMinimum",type=float,default=0.,required=False)

parser.add_argument("--rMaximum",type=float,default=10.,required=False)

parser.add_argument("--dMinimum",type=float,default=0.,required=False)

parser.add_argument("--dMaximum",type=float,default=0.1,required=False)

parser.add_argument("--iterMaximum",type=int,default=1000,required=False)

parser.add_argument("--seed",type=int,default=2020,required=False)

parser.add_argument("--averageHalfWidth",type=int,default=0,required=False)

parser.add_argument("--resetDonTmax",action="store_true",default=False,required=False)

parser.add_argument("experimentResultFiles",nargs='+')

args = parser.parse_args()

timeCutoff	= args.timeCutoff

rMinimum	= args.rMinimum

rMaximum	= args.rMaximum

dMinimum	= args.dMinimum

dMaximum	= args.dMaximum

iterMaximum	= args.iterMaximum

seed	= args.seed

averageHalfWidth	= args.averageHalfWidth

resetDonTmax	= args.resetDonTmax

experimentResultFiles = args.experimentResultFiles

np.random.seed(seed)

modelFitColumns = ['Date',

'Bacteria',

'Wavelength',

'Carbon Source',

'Concentration',

'Replicate',

'Plate',

'Well Row',

'Well Column',

'Bacteria Present',

'r',

'd',

'tmax',

'ymax',

'Tail Slope Std',

'Tlag',

'r*ymax',

'Objective Function',

'Message']

resultsDF = pandas.DataFrame(list(),columns=modelFitColumns)

resultsDF.to_csv("modelFitResults.csv",index=False)

for experimentResultFile in experimentResultFiles:

print("Processing Results in %s" % (experimentResultFile))

results = fitModelToExperiment(experimentResultFile,

timeCutoff,

resetDonTmax,

averageHalfWidth,

rMinimum,rMaximum,

dMinimum,dMaximum,

iterMaximum)

resultsDF = pandas.DataFrame(list(results),columns=modelFitColumns)

resultsDF.to_csv("modelFitResults.csv",index=False,header=False,mode='a')
